# Cerebellar activation in human placebo analgesia: Bridging findings from mice to humans

**DOI:** 10.64898/2026.04.07.717067

**Authors:** Zhaoxing Wei, Tamas Spisak, Dagmar Timmann, Grégory Scherrer, Ulrike Bingel, Tor D. Wager, the Placebo Imaging Consortium

## Abstract

Placebo analgesia has traditionally been explained by top–down cortical regulation of brainstem and spinal pathways. Recent circuit-level work in animal models identified a rostral anterior cingulate–pontine–cerebellar pathway that contributes to expectation-based analgesia, implicating cerebellum circuits in placebo effects^1^. Building on these findings, we examined pontine and cerebellar contributions within a large individual-participant meta-analysis of human neuroimaging studies of placebo analgesia^2^ (*n* = 603). We found that the effects of human pain and placebo converge in cerebellar territories embedded in higher-order cognitive^3,4^ and action-mode networks^5^. These regions exhibit placebo-induced anticipatory increases and reduced responses during painful stimulation, which correlate with the magnitude of placebo analgesia, consistent with predictive configuration of the system. Pontine responses also correlate with individual differences in placebo analgesia. In independent Human Connectome Project data (*n* = 820), pontine activity is functionally connected with cingulate and cerebellar regions implicated in placebo analgesia. Together, these findings support a model in which expectation effects are implemented via predictive configuration of a cortico–pontine–cerebellar system.

## Main

Placebo analgesia, the reduction of pain through expectation, has long been attributed to altered cortical responses during pain and top–down engagement of descending control through brainstem–spinal pathways^6–12^. Recent work broadened this framework by identifying a cortico– pontine–cerebellar pathway in which mice rostral anterior cingulate cortex (rACC) projections to a pontine nucleus (Pn), and then from this nucleus to the cerebellum (rACC→Pn→Cb), are essential for behaviorally conditioned analgesia^1^. In this paradigm, mice that associated a chamber with pain relief showed opioid-dependent reductions in pain behaviors, accompanied by a sharp increase in rACC→Pn activity during the transition into the relief-associated chamber. Optogenetic inhibition of rACC→Pn or Pn→Cb abolished placebo-like analgesia, whereas activation induced analgesia without conditioning. Importantly, cerebellar Purkinje cells displayed anticipatory activity, implicating cerebellar circuits in expectation-driven pain modulation^1^.

Although cerebellar activations and deactivations are frequently observed in human pain and placebo imaging^2,13–15^, they have often been dismissed as motor-related epiphenomena^16^. However, studies across species point to a direct role in modulating spinal nociceptive input and pain. The cerebellum receives input from nociceptive pathways^15,17^, and electrical stimulation of the cerebellar hemispheres reduces nociceptive responses^15,18^. Cerebellar lesions reduce early somatosensory evoked potentials (N24/P24) contralateral to the lesion^19^ and increase pain^15^. Electrical inhibition of cerebellar circuits in humans can increase pain and noxious evoked potentials without affecting motor evoked potentials^20^.

In addition, many cerebellar territories participate in higher-order cognitive networks^21^, and cerebro-cerebellar loops link prefrontal, limbic, affective, and cognitive systems^22–25^. For example, lobule VI and Crus I/II are associated with cortical ‘Salience/Ventral Attention’ and ‘Control’ networks^4^, with cognitive ‘Action’, ‘Demand’, and ‘Social-Linguistic-Spatial’ functional domains^3^, and with the Action-Mode Network (AMN)^5^ **(Fig. 1a–c)**. Lobule VI and Crus I/II are also linked to prediction and expectation, including mismatch and associative learning, for example, cerebellar prediction and prediction-error signals in fear conditioning and extinction^26–28^. Remarkably, Chen et al. found that Pn neurons that receive projections from rACC neurons activated during pain relief expectation precisely project to lobule VI and Crus I/II^1^. Together, these findings suggest a new neural basis for placebo analgesia grounded in prediction-based modulation of pain and implemented in cortico-pontine-cerebellar circuits.

**Fig. 1.**
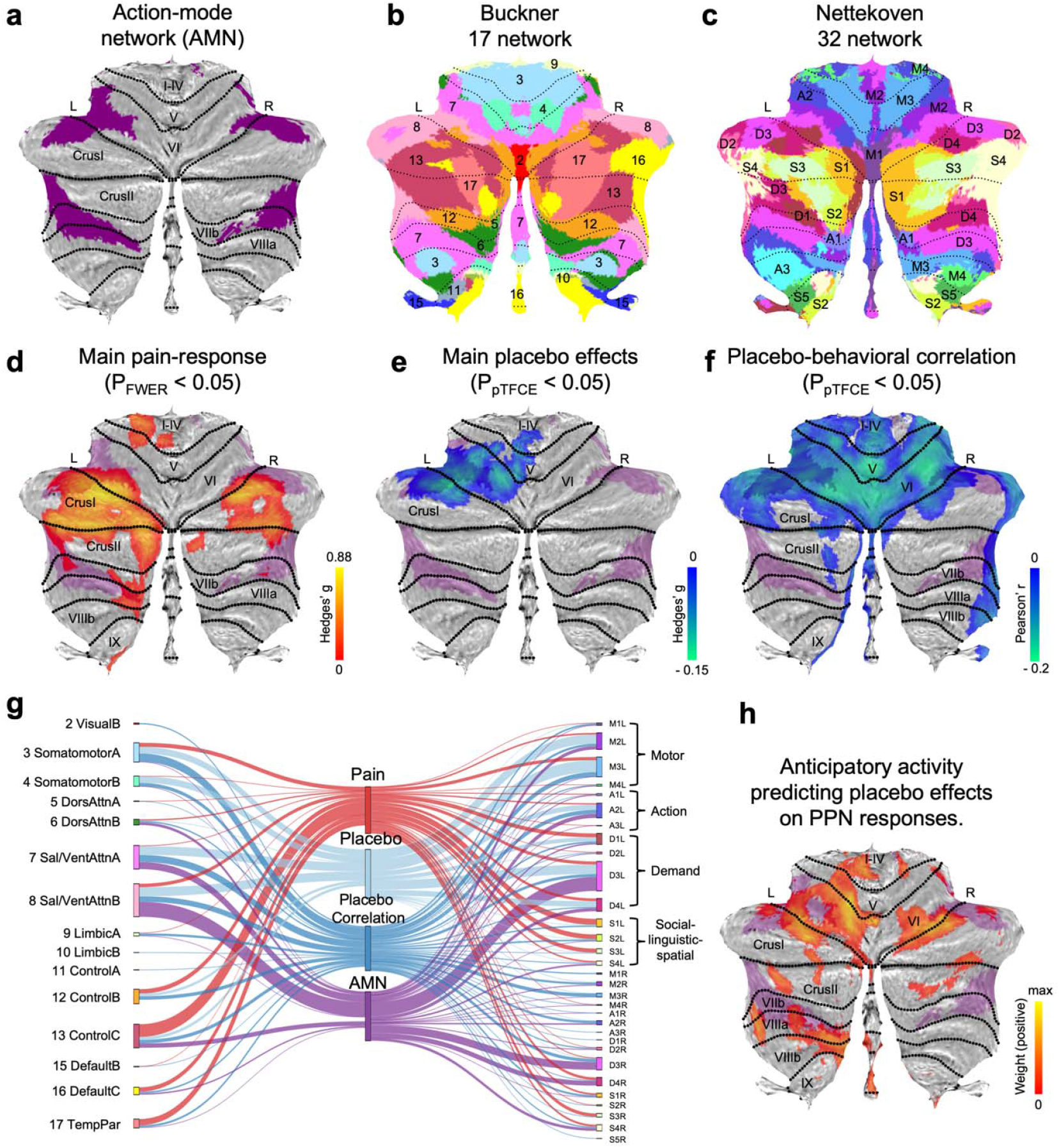
Cerebellar parcellations and meta-analytic maps of pain and placebo analgesia. a–c. Reference parcellations. **a**, Cerebellar flatmap of the Action-Mode Network (AMN)^5^. **b**, Resting-state functional parcellation of the cerebellum from the Buckner et al. 17-network atlas^4^. Each color/number denotes a cortical network represented in cerebellar territories. **c**, Medium-grained 32-network parcellation^3^ grouped into four functional domains: action (A), demand (D), motor (M), and social–linguistic–spatial (S). **d–f. Meta-analytic maps. d**, Pain-evoked random-effects response (Pain > Baseline, *P*_FWE_ < 0.05). Red-yellow colors indicate effect size (Hedges’ *g*). **e**, Significant placebo-related random-effects responses (Hedges’ *g, P*_pTFCE_ < 0.05). Blue-green colors indicate reduced pain-evoked activity in placebo condition relative to control. **f**, Significant correlations between placebo-related activity changes (*Pain*_placebo_ − *Pain*_control_) and placebo analgesia (*Pain*_control_ − *Pain*_placebo_) (Pearson’s *r*; *P*_pTFCE_ < 0.05). Blue–green colors (negative correlations) indicate greater cerebellar decreases with greater placebo analgesia. **g**, The river plot visualizes the correspondence between the activation maps and the AMN across the Buckner 17-network^4^ (left) and the Nettekoven 32-network^3^ (right). Ribbon width reflects Dice similarity. ‘Placebo Correlation’ is the brain placebo analgesia correlation map. **h**, Anticipatory LASSO-PCR weights predicting placebo-induced reductions in pain-processing network (PPN) responses^14^. Red–yellow colors represent positive voxel weights (0 to 7.86 × 10□□). Maps are shown on the SUIT cerebellar flatmap; panels **(d**–**f, h)** are additionally overlaid with the AMN mask (purple). Sample size per voxel: **(d, e)** *n* = 543–603 from 17–20 studies; **(f)**: *n* = 384–460 from 15–18 studies; **(h)**: *n* = 47 from 2 studies. L = left; R = right; I–IV to IX, cerebellar lobules. FWE = family-wise error; pTFCE = probabilistic threshold-free cluster enhancement.

To provide a detailed characterization of pain and placebo effects in the cerebellum, we reanalyzed participant-level data from 20 fMRI studies of placebo analgesia (*n* = 603)^2^, examining the effects of: (i) painful stimulation, (ii) placebo treatment, and (iii) brain–behavior correlations with individual differences in placebo analgesia. In addition, we examined placebo-predictive maps derived from anticipatory-phase fMRI data before painful stimulation in a subset of these studies (*n* = 47). Cerebellar activations were localized and compared across multiple functional parcellations **(Fig. 1a–c)**. Detailed methodological information is provided in the **Supplementary Information**.

Both pain and placebo robustly influenced cerebellar activity, chiefly in regions associated with higher-order control of action and cognition rather than lower-level somatomotor function. Painful stimulation elicited robust cerebellar activity increases broadly associated with higher-order action control and socio-linguistic processes, centered in Crus I/II and lobule VI **(Fig. 1d; Supplementary Fig. 1a)**. This included dorsal AMN regions thought to be recruited during externally focused, action-oriented brain states, and zones associated with multiple resting-state networks, primarily ‘Control’, ‘Temporal-Parietal’ (TempPar), and ‘Default Mode’ as defined by Buckner et al.^4^, as well as zones associated with ‘Social-Linguistic-Spatial’, and ‘Demand’ networks according to Nettekoven et al.^3^ **(Fig. 1g)**.

Placebo treatment, on the other hand, reduced pain-related activity in Crus I and lobule VI, extending to anterior lobules I–IV and V, mainly in the left hemisphere (**Fig. 1e; Supplementary Fig. 1b**). Brain–behavior correlations in placebo analgesia were widespread across both cerebellar hemispheres, spanning lobules I-IV, V, VI, Crus I/II, and the vermis **(Fig. 1f; Supplementary Fig. 1c)**. The regions showing reduced pain responses under placebo were largely contained within the broader zones associated with behavioral analgesia, spanning Buckner et al.’s ‘Somatomotor’ and ‘Salience/Ventral-Attention’ and Nettekoven et al.’s ‘Motor’, ‘Action’, and ‘Demand’ networks. Both pain and placebo overlapped with dorsal AMN regions in Crus I/lobule VI (purple in **Fig. 1a and d-h**). AMN, in turn, overlapped most strongly with ‘Salience/Ventral-Attention’ and ‘Demand’ networks **(Fig. 1g)**.

While the meta-analytic results above characterize cerebellar responses during painful stimulation, Chen et al.^1^ highlight prominent anticipatory increases in cerebellar Purkinje cells. To align our human findings with this anticipatory component, we examined anticipatory placebo-predictive maps before painful stimulation (*n* = 47)^14^. Cerebellar regions showing positive multivariate predictive weights and positive univariate associations with placebo analgesia overlapped extensively with cerebellar regions identified in the large-scale meta-analysis, with prominent overlap in territories associated with the AMN **(Fig. 1h; Supplementary Fig. 2 a-b, Fig. 3)**. Together, these results suggest that cerebellar regions associated with human placebo analgesia exhibit anticipatory increases and pain-phase reductions.

Inspired by Chen et al.’s identification of a placebo-related rACC→Pn→Cb pathway^1^ and the fact that substantial input to Crus I/lobule VI comes from Pn, we examined placebo-related activity in the pons and ACC, along with resting-state functional connectivity between these regions in independent data from the Human Connectome Project. This analysis identified an underappreciated pontine contribution to placebo effects: individuals with stronger placebo analgesia showed larger placebo-related decreases during pain in the pontine nucleus (Pn), located in the basilar part of the pons (*n* = 460, **Fig. 2a)**. The area correlated with placebo analgesia was spatially distinct from the pontine tegmentum (PnTg) **(Supplementary Fig. 1)**, which contains several nuclei involved in descending pain modulation, including the locus coeruleus. We did not find pain or placebo main effects in Pn. To further validate the role of Pn, we examined its resting-state functional connectivity using the Pn as a seed (Human Connectome Project S900, *n* = 820). The strongest connectivity with Pn was found in the anterior cingulate zone of the mid-cingulate cortex (aMCC), medial thalamus, posterior insula, and precuneus, regions associated with pain construction and modulation by placebo and other contextual factors, as well as in the cerebellum, particularly Crus I and lobule VI, paralleling the territories innervated in rodents. The cerebellar results overlapped with pain, placebo, and anticipatory conjunction regions shown in **Fig. 1** and with AMN **(Fig. 2b; Supplementary Fig. 3b and Fig. 5**). These results delineate a candidate circuit for pain construction and position ponto-cerebellar pathways as a key hub.

**Fig. 2.**
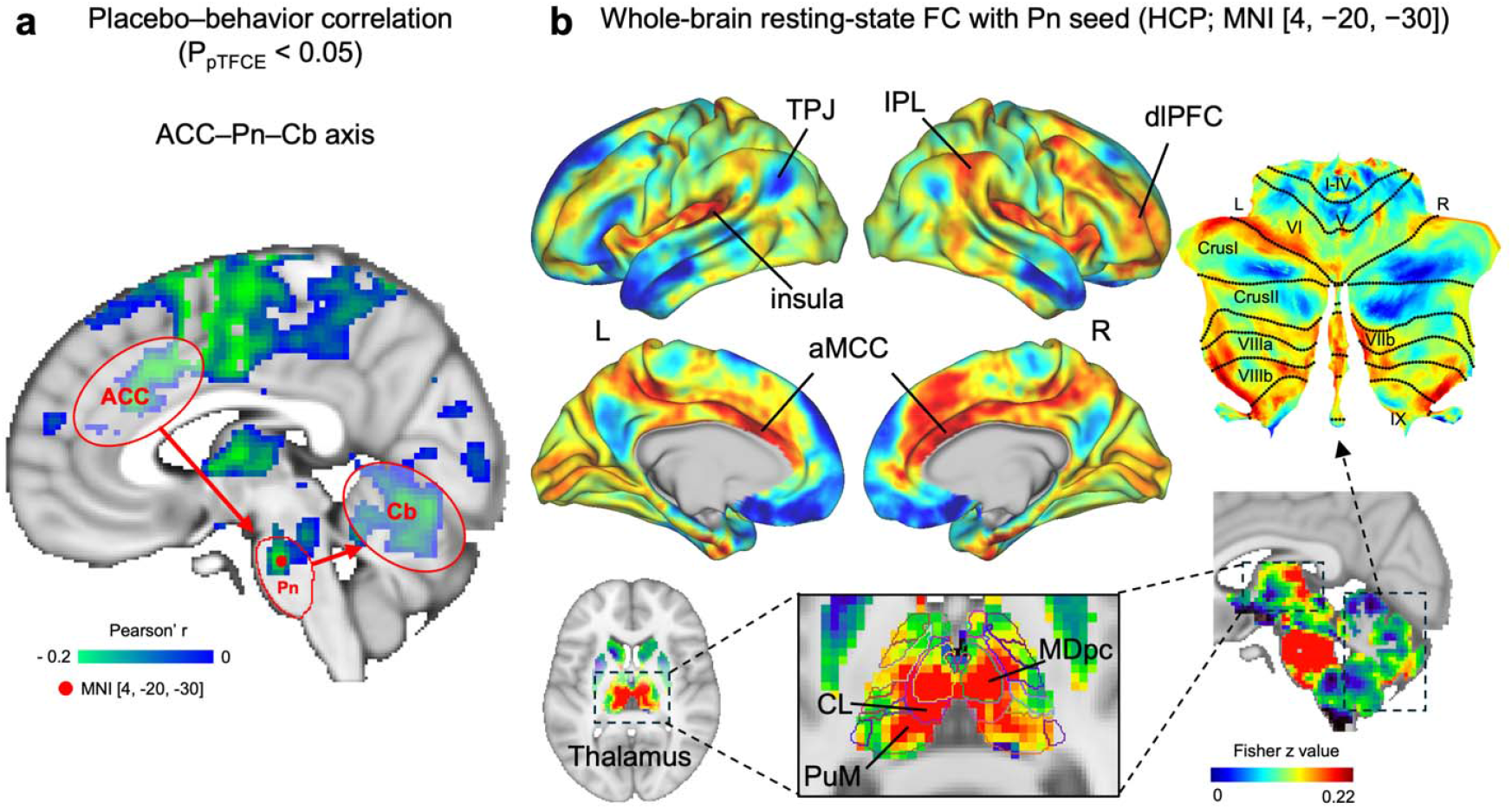
Cortico–pontine–cerebellar axis of placebo analgesia. **a**, Significant brainplacebo analgesia correlations (*P*_pTFCE_ < 0.05), as in **Fig. 1f**. Blue–green colors (negative correlations, *P*_pTFCE_ < 0.05) indicate greater cerebellar decreases with greater placebo analgesia. Significant correlations are found in all elements of the anterior cingulate cortex (ACC)–pontine nucleus (Pn)–cerebellum (Cb) pathway identified in Chen et al^1^. The red dot marks the peak voxel within the Pn (Allen Human References Atlas, 2020, MNI space [4, 20, -30]). b, Group-level whole- brain resting-state functional connectivity seeded at the peak voxel within the Pn (red dot in Figure 2a) was derived from the HCP S900 group connectome (n = 820; MSMAll-aligned, Fisher z-transformed correlation matrix) and visualized on the HCP S1200 MSMAll fs_LR 32k cortical surface, subcortical volume and the SUIT cerebellar flatmap. Warmer colors indicate stronger positive intrinsic connectivity. The strongest correlations in cortex were in the anterior mid-cingulate cortex (aMCC), among the regions most frequently affected in placebo analgesia^8^, as well as the anterior and posterior insula, dorsolateral prefrontal cortex (dlPFC), and inferior parietal lobule (IPL). Subcortically, connectivity encompassed thalamic nuclei, including the mediodorsal parvicellular (MDpc), central lateral (CL), and medial pulvinar (PuM) nuclei (Morel thalamic atlas). In the cerebellum, connectivity was distributed over lobule VI, Crus I, and VIIb/VIIIa/VIIIb, overlapping the cerebellar territories associated with pain, placebo, and anticipatory effects. Together, this intrinsic connectivity pattern is consistent with a cortico– pontine–cerebellar network corresponding to panel (a).

## Discussion

Integrating evidence from human and animal studies, our findings suggest a temporally biphasic model of cerebellar involvement in placebo analgesia, in which cerebellar circuits engaged during pain anticipation preconfigure the system and subsequently manifest as reduced pain-related activity and lower subjective pain during stimulation. This temporal organization is consistent with a predictive computational framework in which cerebellar circuits generate expectations about forthcoming states^29–31^. Within this framework, anticipatory cerebellar engagement in placebo analgesia may reflect the integration of contextual and appraisal-related signals originating in prefrontal and cingulate regions, relayed via pontine nuclei to posterolateral cerebellar territories^1^. This predictive configuration, established prior to nociceptive input, provides an account of how subsequent attenuation of pain-period responses can emerge without requiring direct sensory suppression at the outset^6,32^.

Consistent with this interpretation, circuit-level findings in animal models^1^ demonstrate that during contextual transitions into an analgesia-associated environment, activity in the rACC→Pn pathway increased markedly, accompanied by anticipatory firing of cerebellar Purkinje cells. Correspondingly, in humans, anticipatory increases predictive of placebo analgesia are localized predominantly to posterolateral cerebellar regions, including lobule VI and Crus I/II, whereas during placebo analgesia, these same regions exhibited reduced pain-related responses. Engagement of overlapping circuits at different time points but in opposite directions provides a unified framework reconciling anticipatory increases with placebo-related decreases in cerebellar activity.

An important next step will be to determine whether decreased activity in response to noxious stimuli is observed in animal models of conditioned analgesia. Establishing whether the rACC→Pn→Cb pathway exhibits modulation during noxious stimulation itself will be critical for clarifying whether it directly contributes to sensory pain processing or primarily supports anticipatory modulation. A related question concerns whether this circuit functions mainly to modulate ascending nociceptive afferents in the spinal dorsal horn or instead chiefly confers central nervous system control over pain behaviors and associated action plans. Furthermore, it will be important to establish whether cerebellar involvement in placebo analgesia is restricted to strongly conditioned paradigms or also extends to analgesia driven by higher cognition, beliefs, or social context.

The cerebellar territories in Crus I and lobule VI, which we find to be central to placebo analgesia – and connected with pontine and cingulate regions – have been associated with contextual control and schema-based control over actions, and correspondingly with mid-dorsolateral and anterior prefrontal cortices^29^. Anterior cingulate is likewise thought to play a direct role in pain-related action policy and only a more indirect role in the regulation of nociception^5,33^. A current understanding of this circuit suggests that placebo analgesia may act not only through modulation of sensory afferent signals but also via reconfiguration of action-planning and control systems that shape readiness for upcoming sensory events^12,34^. At a systems level, disconnection in superior cerebellar regions overlapping with placebo-related and AMN regions was associated with a chronic nociplastic pain in the UK Biobank^35^ (**Supplementary Fig. 2d**). Disruption of this circuit may thus contribute to maladaptive pain persistence. This systems-level vulnerability aligns with evidence that sensory expectations can directly shape population-level dynamics within motor circuits^36^, providing a mechanistic basis for cerebellar and motor-network engagement under placebo conditions. By this view, recruitment of somatomotor cerebellar territories–like AMN and inter-effector regions in M1–may reflect anticipatory preparation in motor and action-planning circuits ^37,38^ based on potential future harm.

An open question is the relationship between control of pain and action in the cerebellum. Pain and action may be mutually inhibitory, and the cerebellar territories involved in AMN and action control may be activated as part of an action planning-based suppression of pain. Alternatively, placebo effects on the cerebellum may relate more directly to higher-level cognitive processes implemented in cerebello-prefrontal circuits. Finally, cerebellar activity may relate more directly to action inhibition in fMRI settings, in which participants experience painful stimuli without withdrawing. However, the direct modulation of pain and early somatosensory potentials^15,18–20^ argues against a purely motor or motor-inhibition role.

Future cross-species work combining conditioning paradigms with targeted circuit manipulations will be essential to dissociate predictive and sensory contributions and to determine how cerebellar outputs influence downstream cortical and brainstem pain-control regions. Non-human studies could investigate how this circuit regulates nociceptive activity in descending (periaqueductal gray (PAG), rostroventromedial medulla (RVM), dorsal horn neurons) and ascending nociceptive pathways. Human studies should systematically incorporate the cerebellum into human pain neuroimaging, including full cerebellar coverage, network-level analyses, and explicit testing of cortical–brainstem-cerebellar connectivity. Establishing causal mechanisms in humans will require complementary approaches such as cerebellar lesion studies^39^ and transient modulation of cerebellar excitability^40^. In this context, the intrinsic connectivity pattern identified here **(Fig. 2b)** provides a roadmap for such studies, highlighting cortical, thalamic, pontine, and cerebellar nodes that participate in this circuit and could be targeted by causal manipulations. Coupled with those of Chen et al., our findings motivate a reassessment of prevailing corticocentric models of cognitive and affective control and suggest new neuromodulatory targets within the cortico-pontine-cerebellar system for predictive control over pain.

## Acknowledgements

T.D.W. was supported by the National Institutes of Health (NIH) / National Institute of Mental Health (NIMH) under grant MH R37076136. T.S., D.T., and U.B. were supported by the German Research Foundation (DFG; TRR 289, Project-ID 422744262).

