## Supplementary Information for "Cerebellar activation in human placebo analgesia: Bridging findings from mice to humans"

**# The Placebo Imaging Consortium Authors**

Lauren Atlas^4,5,6^, Fabrizio Benedetti^7,8^, Ulrike Bingel^2^, Christian Büchel^9^, Jae Chan Choi^10,11,12^, Luana Colloca^13^, Davide Duzzi^14^, Falk Eippert^15^, Dan‑Mikael Ellingsen^16,17^, Sigrid Elsenbruch^2,18^, Ted J. Kaptchuk^19^, Simon S. Kessner^20^, Irving Kirsch^19^, Jian Kong^21^, Claus Lamm^22^, Siri Leknes^23,24^, Fausta Lui^25^, Alexa Müllner‑Huber^22^, Carlo A. Porro^25^, Markus Rütgen^22^, Lieven A. Schenk^9^, Tamás Spisák^2^, Irene Tracey^26^, Tor D. Wager^1^, Fadel Zeidan^27^, Matthias Zunhammer^2^

**Supplementary Figures**

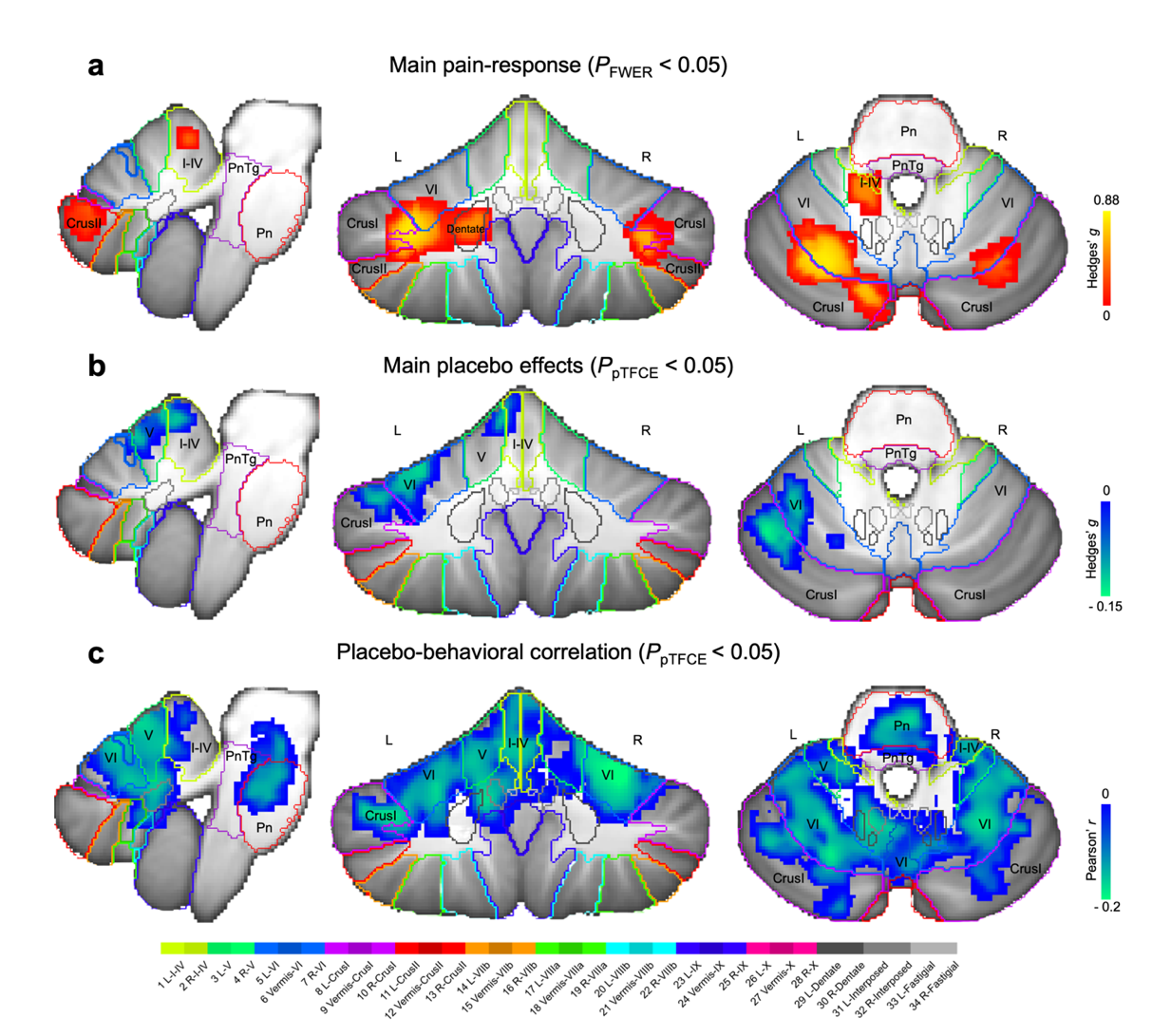

**Supplementary Fig. 1. Cerebellar activations shown in SUIT volume space (corresponding to Fig. 1d–f). a,** Main pain-evoked response. Random-effects group activation for *pain > baseline*, identified by non-parametric permutation testing and corrected for family-wise error rate (*P*_FWE_ < 0.05, two-sided). Red–yellow colors indicate effect size (Hedges’ *g*). **b,** Main placebo effects. Significant placebo-related random-effects responses (*P*_pTFCE_ < 0.05; Hedges’ *g*). Blue–green colors indicate larger negative effects (reduced pain-evoked activity during placebo). **c,** Placebo–behavioral correlation. Cerebellar activations showing significant correlations between individual behavioral placebo analgesia (*Pain*_control_ − *Pain*_placebo_) and placebo-related neural changes (*Pain*_placebo_ − *Pain*_control_) (Pearson’s *r*; *P*_pTFCE_ < 0.05). Blue–green colors indicate stronger correlations between cerebellar activation and the magnitude of behavioral analgesia. Left panel: sagittal view (left hemisphere); middle: axial (horizontal) view; right: coronal view. The pontine nucleus (Pn) and pontine tegmentum (PnTg) are delineated within the pontine region of the brainstem for anatomical reference. Colored outlines in the cerebellum indicate lobular boundaries from the Diedrichsen 34-lobular atlas^6^ in SUIT volume space. The color bar is shown at the bottom. L, left hemisphere; R, right hemisphere.

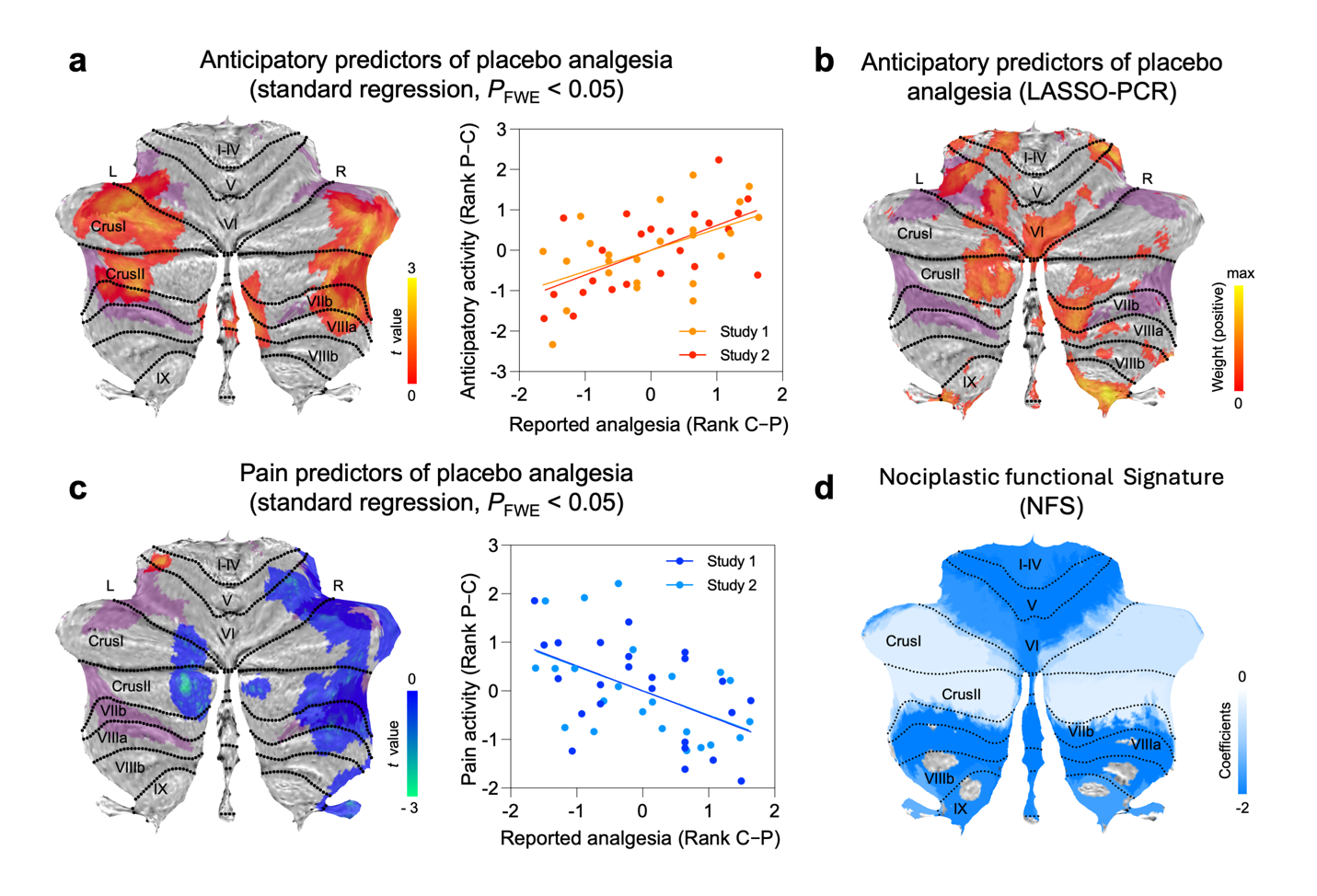

**Supplementary Fig. 2. Cerebellar predictors of placebo analgesia and nociplastic functional connectivity signature. a, c,** Cerebellar predictors of placebo analgesia during the anticipatory phase (**a**) and pain phase (**c**)^5^, assessed using behavioral pain ratings. Voxelwise regression maps identify cerebellar regions in which activity during anticipatory (*Antic*_placebo_ − *Antic*_control_) and pain phase (*Pain_p_*_lacebo_ − *Pain*_control_) predicts individual differences in placebo analgesia (*Pain*_control_− *Pain*_placebo_). Warm colors in (**a**) indicate positive associations, whereas cool colors in (**c**) indicate negative associations. Scatter plots show mean cerebellar cluster activity change versus reported analgesia (within-study ranked values), with separate fit lines for Study 1 and Study 2. **b,** Multivariate anticipatory predictors of placebo analgesia measured by behavioral pain ratings^5^. Voxel weight maps indicating anticipatory neural activity predictive of individual differences in behavioral placebo analgesia. Red–yellow colors depict positive predictive weights, illustrating voxels in which higher anticipatory responses predict stronger behavioral analgesia. **d,** Visualization of the nociplastic functional signature (NFS) connectivity pattern, thresholded to emphasize the top 25% of structure coefficients, which denote the summed dynamic conditional correlations across brain parcels. All maps are displayed on cerebellar SUIT flatmaps, with the AMN overlaid in purple in panels a–c for interpretability.

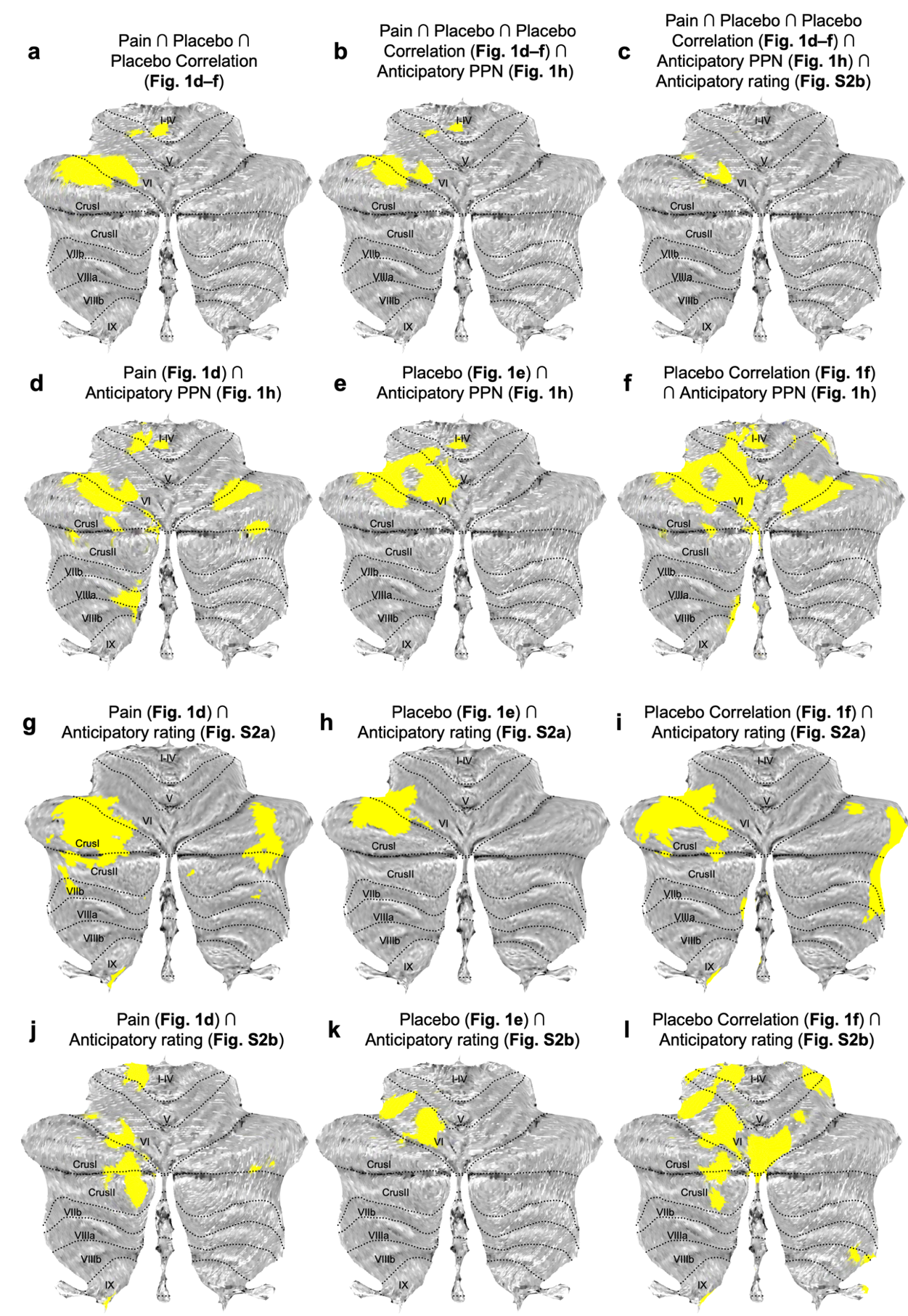

**Supplementary Fig. 3. Spatial overlap between pain, placebo, behavioral correlation, and anticipatory cerebellar masks. a,** Three-way overlap between pain-evoked increases, placebo-related decreases, and placebo–behavioral correlations derived from the meta-analysis (corresponding to **Fig. 1d–f**). **b,** Overlap between the anticipatory activity predicting placebo effects on pain-processing network (PPN) responses and the three-way meta-analytic overlap shown in (a). **c,** Overlap across all five masks: pain, placebo, behavioral correlation, anticipatory PPN-predictive activity, and anticipatory rating-predictive activity. **d–f,** Overlaps between anticipatory PPN-predictive activity and each meta-analytic component separately: pain-evoked increases (d), placebo-related decreases (e), and placebo–behavioral correlations (f). **g–i,** Overlaps between anticipatory rating-predictive map (univariate) and each meta-analytic map separately: pain-evoked increases (g), placebo-related decreases (h), and placebo–behavioral correlations (i). **j–l,** Overlaps between anticipatory rating-predictive map (multivariate) and each meta-analytic map: pain-evoked increases (j), placebo-related decreases (k), and placebo–behavioral correlations (l). Binary overlap maps are displayed on the SUIT cerebellar flatmap. Yellow regions indicate vertices shared between the indicated masks; non-overlapping vertices are shown in grey. The overlap operator (∩) denotes the spatial intersection.

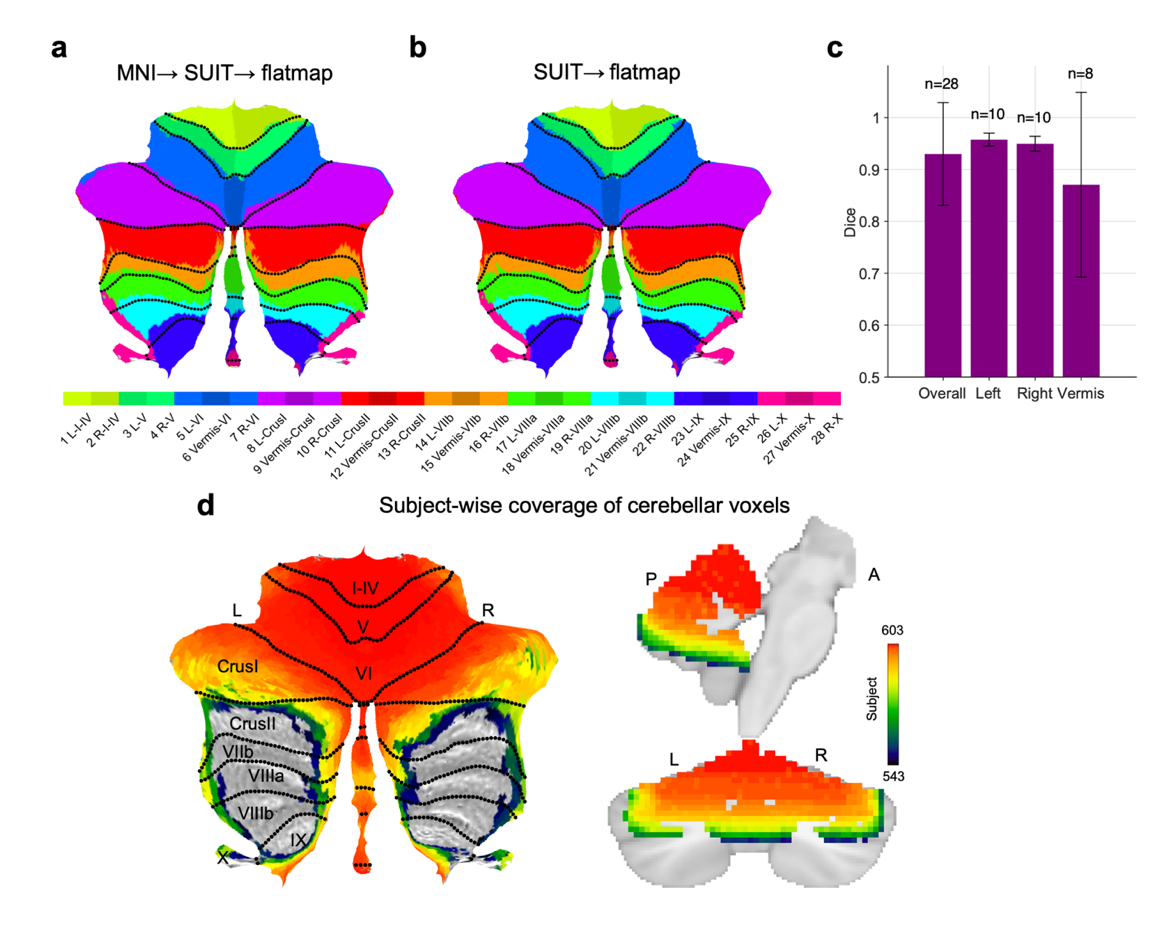

**Supplementary Fig. 4. Registration accuracy on the cerebellar flatmap.** **a,** Diedrichsen 34-region atlas^6^ defined in MNI space and displayed after transformation to SUIT (MNI→SUIT→flatmap). **b,** Native SUIT version of the same atlas displayed on the flatmap (SUIT→flatmap). Colors denote atlas labels; dotted black curves indicate lobular boundaries. **c,** Bar plots show the mean Dice similarity coefficient (±SD) between (a) and (b), summarized for all cerebellar ROIs and by subgroups (Left, Right, Vermis). Dice coefficients are above 0.9 for left and right hemispheres and overall, indicating good-quality mapping from MNI to the cerebellar surface at the macro scale. Error bars indicate SD; n is the number of regions in each subgroup. **d,** Subject-wise coverage of cerebellar voxels of pain and placebo contrast, projected onto the SUIT flatmap (left) and MNI152 space (right). Each voxel encompasses data from 543 to 603 individuals across 17 to 20 independent studies. I–IV to IX, cerebellar lobules. L, left hemisphere; R, right hemisphere.

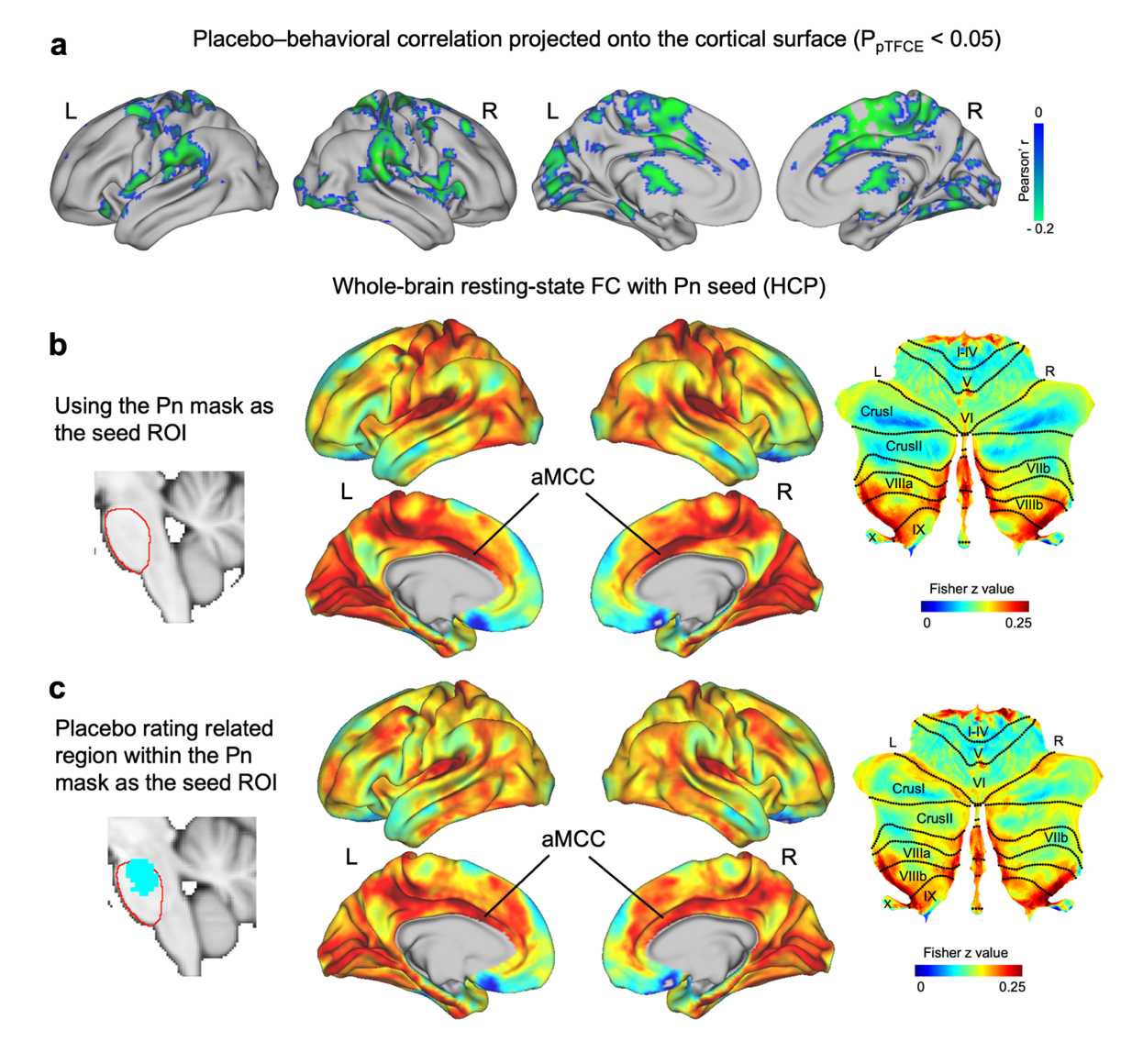

**Supplementary Fig. 5.** **Cortico-cerebellar and pontine connectivity patterns associated with behavioral placebo effects. a,** Cortical projection of placebo–behavioral correlations (*P*_pTFCE_ < 0.05), showing surface-wide associations between behavioral analgesia (*Pain*_control_ – *Pain*_placebo_) and placebo-related activity changes (*Pain*_placebo_ − *Pain*_control_). Green–blue colors indicate stronger positive correlations (Pearson’s *r*). **b,** Whole-brain resting-state functional connectivity map derived from the Human Connectome Project (HCP, *n* = 820) using the pontine nucleus (Pn) as the seed region of interest (ROI; red outline, left). The resulting Fisher-*z*–transformed map shows intrinsic connectivity between the Pn and distributed cortical and cerebellar regions, with the strongest coupling observed in the anterior mid-cingulate cortex (aMCC) and lobules VI, Crus I/II, and VIII. **c,** Functional connectivity map generated using the placebo-rating–related subregion within the Pn as the seed ROI (cyan area, left). The resulting pattern largely overlaps with the whole Pn seed map, highlighting consistent connectivity between pontine, cerebellar, and cingulate regions that form a potential brainstem–cerebellar–cortical pathway mediating placebo analgesia. Color bar: Fisher-*z* values (0–0.25). L/R, left/right hemisphere; I–IV to IX, cerebellar lobules.

**Supplementary Methods**

**Meta-analytic dataset and statistical inference**

**Data acquisition**. We analyzed group-level statistical maps from a prior individual-participant data meta-analysis of 20 published studies^1^. A detailed description of the data collection procedures can be found in Zunhammer et al^2^. The outcome definitions, effect-size standardization, and voxelwise statistical inference follow the original meta-analysis. We summarize them here for completeness and did not re-estimate whole-brain statistics. The corresponding 3D volumes are publicly available at <https://osf.io/n9mb3/>.

**Outcomes and comparisons**. We evaluated three voxelwise outcomes from the prior meta-analysis. All voxelwise statistical maps were aggregated across studies using a random-effects model, which treats each study’s effect as a sample from a broader population distribution and accounts for between-study heterogeneity.

1. Main pain stimulation effects (Pain > Baseline). Mean pain-evoked response relative to baseline. When both placebo and control sessions were available within a study, the two were averaged to obtain a single pain-versus-baseline estimate, accommodating studies that reported only pooled main effects.
2. Placebo treatment effect (*Pain*_placebo_ − *Pain*_control_). The analysis quantified the difference in pain-evoked activity between placebo and control conditions, indexing how placebo treatment modulates pain-related activation.
3. Brain–behavior correlation in placebo analgesia. In within-subject placebo designs, the analysis computed across-participant correlations between the neural placebo effect and behavioral placebo analgesia. Using (*Pain*_placebo_ − *Pain*_control_) for neural maps and (*Pain*_control_ − *Pain*_placebo_) for behavioral ratings ensures aligned directions of effect.

**Effect-size scaling and brain–behavior correlations.** For the main pain effect and the neural placebo effect, study-wise maps were standardized using each study’s between-subject SD of pain-related activity to obtain Hedges’ g (bias-corrected Cohen’s *d*). For within-subject designs, Hedges’ g_rm (repeated-measures variant) was used. This small-sample–corrected scaling places all studies on a common metric before aggregation.

For the coupling analysis (18 studies; 460 participants), voxelwise Pearson’s r was computed within each study between behavioral placebo analgesia and the neural placebo effect, then transformed using Fisher’s z for across-study averaging and inference (with back-transformation for reporting where appropriate).

**Voxelwise statistical inference.** Voxelwise *P* values were obtained with a nonparametric permutation test of the pseudo-*z* statistic, controlling the family-wise error (FWE) via the maximum-statistic (max-*z*) approach (1,500 permutations; for minimal *p* values, *P* < 0.01, the permutation tail was approximated by a generalized Pareto fit). For the pain main effect, only the unenhanced max-*z* FWE procedure was used; maps were thresholded at two-tailed FWE-corrected α = 0.05 (*P*_FWE_ <0.05). For the placebo main effect and the placebo–rating correlation, probabilistic threshold-free cluster enhancement (pTFCE) was additionally applied, and maps were thresholded at two-tailed *P_p_*_TFCE_<0.05.

**Anticipatory placebo-predictive maps**

To facilitate comparison with the anticipatory neural dynamics described in Chen et al.^3^, we examined voxel-wise predictive maps derived from anticipatory-phase fMRI data from two studies (*n* = 47), originally reported in Wager et al.^4^ and subsequently reanalyzed using LASSO–PCR^5^. These maps represent the only publicly accessible anticipatory-phase data within the 20 published studies^1^. Because the published LASSO–PCR weight maps do not include brainstem voxels, we focus here on positive cerebellar weights to enable direct comparison with cerebellar anticipatory signals reported in Chen et al.^3^. We include two multivariate LASSO–PCR weight maps: (1) Placebo_Predict, which predicts individual differences in placebo analgesia, operationalized as placebo-related reductions in pain ratings, from anticipatory activity (corresponding to Figure 3c in Wager et al.^5^). (2) Placebo_Brain_Predict, which predicts placebo-related changes in an averaged pain-processing network (PPN) response during pain from anticipatory activity (corresponding to Fig. 7b in Wager et al.^5^).

We additionally constructed cerebellar univariate predictor maps from the same pooled dataset using the regression framework of Wager et al.^5^. In both source studies, anticipatory and pain-phase responses were estimated from designs that included a jittered interval between cue onset and pain delivery, permitting phase-specific modeling. In Study 1, a 3-s warning cue was followed by a jittered anticipatory interval of 3-12 s before a 6-s shock stimulus. In Study 2, a 1-s cue (“Get ready!”) was followed by a jittered anticipatory interval of 1-16 s (mean = 9.77 s) before thermal pain delivery. For anticipatory and pain-phase analyses, input contrasts were control-minus-placebo maps. Within each study (study 1, n = 24; study 2, n = 23), voxel values were rank-transformed across subjects at each voxel. Behavioral placebo analgesia (Pain_control_ − Pain_placebo_) was likewise rank-transformed within study, with the strongest analgesia assigned rank 1, following the original implementation of Wager et al.^5^. Rank-transformed images were then entered into robust voxelwise regression, with placebo analgesia rank as the predictor of interest and study, testing order, and their interaction included as covariates.

**Cerebellar analysis and visualization**

All analyses and flatmap visualizations were mainly performed in MATLAB R2024b using CANlab Core Tools neuroimaging analysis toolbox (<https://github.com/canlab/CanlabCore>) and the SUIT cerebellar toolbox for SPM12 (<https://github.com/jdiedrichsen/suit>). Volume renderings were generated with FSLeyes. Surface visualization and related analysis were performed using Connectome Workbench (v2.0.1; Human Connectome Project).

**Atlas and region of interest (ROI).**

To compare and localize condition-specific statistical maps within the cerebellum, we used two atlases from the Diedrichsen Lab cerebellar atlas collection (<https://github.com/DiedrichsenLab/cerebellar_atlases>), plus a hypothesis-driven Action mode network mask; all resources were in Spatially Unbiased Infratentorial Template (SUIT) flatmap space.

**Anatomical reference.** Diedrichsen 34-region atlas^6^: lobular parcellation (I–X with subdivisions) plus deep nuclei (ROIs 29–34); used to report effects in conventional lobular terms.

**Functional reference.** We used two cerebellar functional atlases from the Diedrichsen Lab collection. The **Buckner 17-network parcellation**^7^ subdivides seven canonical large-scale cortical networks into 17 connectivity-defined cerebellar subsystems**. The Nettekoven 32-network** asymmetric parcellation^8^ provides a fine-grained task- and connectivity-derived functional atlas comprising 32 cerebellar regions organized into four domains: motor (M1–M4), action (A1–A3), demand (D1–D4), and social–linguistic–spatial (S1–S5).

**Action-Mode Network (AMN)**^9^. We used a SUIT-flatmap mask indexing cerebellar territories for sustained executive action control (task-set maintenance, performance monitoring, salience-guided engagement), closely related to the cingulo-opercular system.

**Cerebellar volume and flatmap visualization**

All statistical maps were projected onto the SUIT cerebellar flatmap^10,11^ for visualization and spatial statistics**.**

Statistical maps from the Zunhammer et al.^1^ meta-analysis (pain, placebo, and placebo–rating correlation; see their Figs. 1a, 2a, and 3a) and anticipatory placebo-predictive maps^5^ were first resliced from MNI152NLin6Asym to SUIT space using *suit_reslice_dartel*. For each contrast, a brainstem-cerebellum-only SUIT-space volume was obtained. For 3D volume views **(Supplementary Fig. 1)**, statistical maps were overlaid on the SUIT T1-weighted template with outlines of the 34 cerebellar anatomical ROIs^6^ using *fsleyes*.

SUIT-space volumes were then projected to the 2D cerebellar flatmap with *suit_map2surf* using thickness-averaged sampling (mean across cortical depth) and rendered with *suit_plotflatmap*; positive and negative effects were displayed separately with divergent color maps **(Fig. 1d-f, h; Supplementary Fig. 2a-c)**.

The cerebellar univariate predictor maps (n = 47) were restricted to the cerebellum using the Diedrichsen (2009) mask, resampled to MNI152NLin6Asym 2-mm space. Cluster-wise family-wise error (FWE) correction was applied within the cerebellar mask using cluster-extent thresholding with a cluster-forming threshold of P < 0.05. Significant maps were resliced into SUIT space and visualized on cerebellar flatmaps with lobular boundaries overlaid.

For visualization, subject-level cerebellar values were extracted as the mean rank-transformed signal within significant clusters, z-scored within study, and plotted against within-study z-scored behavioral ranks, with separate regression lines for each study (**Supplementary Fig. 2**).

The NFS map was first resampled to the MNI152NLin6Asym 1-mm template grid using SPM12 with nearest-neighbor interpolation to minimize interpolation-induced smoothing and preserve the sparsity of thresholded weights. The resampled image was then transformed from MNI152NLin6Asym space to SUIT space using *suit_reslice_dartel*, with a SUIT cerebellar mask applied to restrict analyses to cerebellar voxels. The resulting SUIT-space image was projected onto the cerebellar surface using *suit_map2surf* and visualized using *suit_plotflatmap*. For visualization, positive values were excluded by masking the map at zero, and only negative coefficients were displayed, consistent with the directionality of the NFS weights reported in the original study.

To quantify the spatial correspondence between pain-, placebo-, and anticipation-related cerebellar effects, we next performed spatial intersection analyses on the SUIT flatmap. Statistical maps from the Zunhammer et al. meta-analysis (pain, placebo, and placebo–rating correlation) and anticipatory placebo-predictive maps were thresholded and binarized to generate vertex-wise masks. All masks were aligned in SUIT flatmap space, and spatial overlaps were computed using logical conjunction across masks. Resulting overlap maps were visualized on the SUIT flatmap **(Supplementary Fig.** **3)**. The overlap operator (∩) denotes the spatial intersection.

**Cerebellar network overlap analysis**

To quantify spatial correspondence, we performed vertex-wise overlap analyses in SUIT flatmap space^11^ between cerebellar effect maps and functional parcellations. Specifically, three cerebellar statistical maps (pain, placebo, and placebo–behavioral correlation), as well as the AMN map, were each compared against the Buckner 17-network atlas and the Nettekoven 32-network parcellation.

All network atlases were converted into binary ROI masks. To ensure that overlap metrics were computed only in cerebellar regions with empirical data support, all atlas masks were first intersected with a cerebellar coverage mask derived from the meta-analytic pain and placebo contrasts (**Supplementary Fig. 4d**). Overlap analyses were therefore restricted to cerebellar vertices that were both assigned to a given network and covered by the meta-analytic dataset.

For each map–ROI pair, spatial overlap was quantified using the Dice similarity coefficient:

$$Dice\left( A,B \right)= \frac{2 |A\cap B|}{|A|+|B|}$$

where A denotes a binarized cerebellar effect map (or AMN map), and B a binary network ROI, with both masks defined within the coverage-restricted SUIT flatmap space, and |A| and |B| represent the number of vertices within each mask. All overlap results are summarized in **Fig. 1g.**

**Validation analyses**

**Registration accuracy on the cerebellar flatmap.** Because the meta-analytic maps were computed in MNI space but now visualized on the SUIT flatmap, we evaluated the registration fidelity of the transformation chain (MNI→SUIT→ flatmap). We used the Diedrichsen et al.^6^ cerebellar probabilistic atlas (34 lobular regions) defined in both MNI and SUIT space. Each MNI ROI was resliced to SUIT space using suit_reslice_dartel (default interpolation) and binarized at 0.5 to preserve discrete labels. Both MNI→SUIT and native SUIT atlases were projected to the flatmap using suit_map2surf with through-thickness aggregation (@nanmax) and thresholded at >0.5. Registration validation confirmed that the flatmap projections provide accurate macroscale correspondence between MNI-space maps and the cerebellar surface (**Supplementary Fig. 4a-c**). Dice similarity coefficients were computed between corresponding lobular ROIs to quantify projection accuracy. We summarized Dice across all regions and within subsets (left, right, vermis), and visualized group means with SD error bars and sample sizes **(Supplementary Fig.4c)**. The dentate, interposed and fastigial nuclei (labels 29–34) are not represented on the SUIT flatmap and were therefore excluded from the Dice calculation. Across the remaining 28 regions, the MNI→SUIT transformed atlas showed good correspondence with the native SUIT atlas (mean *Dice* = 0.930 ± 0.099; *n* = 28). Agreement was excellent in the left (0.958 ± 0.013; *n* = 10) and right hemispheres (0.950 ± 0.014; *n* = 10), and remained good for vermis regions (0.871 ± 0.178; *n* = 8).

**Subject-wise cerebellar coverage check.** This analysis is based on Supplementary Fig. 2 in Zunhammer et al. ^1^. Group-level maps were first restricted to the probabilistic cerebellar mask using *fslmaths*, then resliced from MNI152NLin6Asym to SUIT space using *suit_reslice_dartel.* The resulting SUIT-space volumes were projected onto the cerebellar flatmap with *suit_map2surf* using through-thickness aggregation (*@nanmax*). Full-sample voxel-wise N maps were processed identically to verify cerebellar coverage, and color bars were scaled to the observed N ranges. Coverage checks showed robust sampling across the cerebellum, with limited coverage in inferior regions (Crus II, VIIb, VIIIa/b, IX), potentially underestimating ventral lobule contributions **(Supplementary Fig. 4d)**.

**Pontine functional connectivity analysis**

We utilized the Human Connectome Project (HCP) S900 group-average dense connectome, a publicly released group-level functional connectivity dataset derived from resting-state fMRI data of 820 participants (R820 group; released December 2015). This dataset is available through the ConnectomeDB portal (<https://db.humanconnectome.org/>). Specifically, we used the file *HCP_S900_820_rfMRI_MSMAll_groupPCA_d4500ROW_zcorr.dconn.nii*, which represents a Fisher-z–transformed grayordinate × grayordinate functional connectivity matrix aligned to the MSMAll-registered fs_LR midthickness 32k surface space (for cortical vertices) and the MNI152NLin6Asym space (for subcortical and cerebellar voxels).

Using this group-average connectome, we examined whole-brain functional connectivity patterns of the pontine nucleus (Pn). A single-voxel seed located in the Pn was defined as the peak voxel of the placebo–behavioral correlation map within the Allen Human Brain Atlas mask (2020, MNI space) **(Fig. 2a)**, identified using *fslstats*. The corresponding row from the HCP dense connectome was extracted to generate the seed-based whole-brain connectivity map, representing Fisher-z–transformed correlations with all 91,282 grayordinates. In addition, both the entire Pn mask and the placebo–rating–related subregion within the Pn were used as complementary ROIs for confirmatory analyses **(Supplementary Fig. 5b,c)**. For each ROI, we identified all subcortical grayordinates contained within the mask. We computed the mean of the corresponding Fisher-z correlation values across these grayordinates within the group dense connectome, yielding an averaged whole-brain connectivity map that represents the mean coupling pattern of the pontine nucleus region.

All resulting maps were visualized on the cortical surface (fs_LR 32k), subcortical volume (MNI152NLin6Asym), and cerebellar surface (SUIT) for comparison with the single-voxel seed results. For quantitative ROI-based analyses, mean connectivity values were further extracted using the CANlab 2024 functional atlas, providing a unified parcellation framework across cortical, subcortical, and cerebellar regions **(Fig. 2a; Supplementary Fig. 5b,c).**
